# High-throughput screening assay for PARP-HPF1 interaction inhibitors to affect DNA damage repair

**DOI:** 10.1101/2023.11.14.566986

**Authors:** Saurabh S. Dhakar, Albert Galera-Prat, Lari Lehtiö

## Abstract

ADP-ribosyltransferases PARP1 and PARP2 play a major role in DNA repair mechanism by detecting the DNA damage and inducing poly-ADP-ribosylation dependent chromatin relaxation and recruitment of repair proteins. Catalytic PARP inhibitors are used as anticancer drugs especially in the case of tumors arising from sensitizing mutations. Recently, a study showed that Histone PARylation Factor (HPF1) forms a joint active site with PARP1/2. The interaction of HPF1 with PARP1/2 alters the automodification site from Aspartate, Glutamate to Serine, which has been shown to be a key ADP-ribosylation event in the context of DNA damage. Therefore disruption of PARP1/2-HPF1 interaction could be an alternative strategy for drug development to block the PARP1/2 activity. In this study, we describe a FRET based high-throughput screening assay to screen inhibitor libraries against PARP-HPF1 interaction. We optimized the conditions for FRET signal and verified the interaction by competing the FRET pair in multiple ways. The assay is robust and easy to automate. Validatory screening showed the robust performance of the assay, and we discovered two compounds, Dimethylacrylshikonin and Alkannin, with µM inhibition potency against PARP1/2-HPF1 interaction. The assay will facilitate the discovery of inhibitors against HPF1-PARP1/2 complex and to develop potentially new effective anticancer agents.

## Introduction

ADP-ribosylation of proteins is a post-translational modification which is involved in regulation of many biological processes including DNA damage repair, transcription, and cell cycle regulation (Gibson & Kraus, 2012). There are 17 ADP-ribosyltransferases (ARTs) in human with homology to Diphtheria toxin (ARTDs) (Lüscher *et al*, 2022). These PARP enzymes and tankyrases regulate a wide range of signaling events and three of them, PARP1-3, are known to recognize DNA lesion sites and activate DNA repair cascades by transferring ADP-ribosyl moiety from NAD^+^ to DNA damage site, histones and to enzyme itself (Lüscher *et al*, 2022; Schreiber *et al*, 2006; Javle & Curtin, 2011; Amé *et al*, 1999). The ADP-ribose moiety is transferred in the form of single unit, mono-ADP ribosyl (MAR), and can be subsequently elongated to form poly-ADP-ribose chains (PAR). The PARylation of histone tail leads to nucleosome remodeling and PARP automodification is a regulatory mechanism for their own activity (De Vos *et al*, 2012). The modification occurs on aspartate, glutamate and serine residues but in presence of Histone PARylation Factor (HPF1), PARP1/2 are more specific and predominantly modify serine residues (Leidecker *et al*, 2016; Bonfiglio *et al*, 2017; Palazzo *et al*, 2018; Crawford *et al*, 2018; Larsen *et al*, 2018). The preference to serine is enabled by HPF1 as it forms a joint active site with PARP1 and PARP2 as demonstrated by a crystal structure of the PARP1/2 catalytic domain – HPF1 complex (Suskiewicz *et al*, 2020; Sun *et al*, 2021).

The role of PARP enzymes in DNA repair and development of small molecule inhibitors gained attention and interest when it was discovered that such catalytic PARP inhibitors have a synthetic lethality with BRCA mutations frequent in e.g. breast cancer (Farmer *et al*, 2005; Bryant *et al*, 2005). Subsequently, multiple PARP inhibitors have been clinically approved for different indications (Li *et al*, 2023). The approved drugs compete with NAD^+^ substrate and thereby block the catalytic site of enzyme. The small molecules are not often very selective (Wahlberg *et al*, 2012) and attempts have been made to generate PARP1 selective inhibitors (Jamal *et al*, 2022) and on the other hand compounds with dual activities (Plummer *et al*, 2020).

An alternative specific strategy to inhibit serine ADP-ribosylation in DNA repair could be to prevent binding of HPF1 to PARP1 and to PARP2. HPF1 initiates PARylation on specific serine residues of histone tails subsequently leading to remodeling of the nucleosome with help of other factors such as ALC1 recognizing the modification (Bacic *et al*, 2021). As HPF1 shares a joint active site with PARP1/2 and plays a key role in increasing enzymatic activities, the inhibition of the interaction between HPF1-PARP would reduce histone PARylation upon DNA damage as single agent or possibly in combination with catalytic PARP inhibitors or DNA damaging agents. In the crystal structure, HPF1 and PARP2 interaction is mediated by a 15 amino acid long helix (261-284) of HPF1 packing against the PARP2 catalytic domain and NAD^+^ binding cleft (Suskiewicz *et al*, 2020) (**Fig. 1a**). Through this, HPF1 and PARP2 collectively form a joint active site. The last two residues of PARP2 C-terminus interact at the back of HPF1 C-terminal domain and the overall interface buried is 1328.6 Å^2^ (Krissinel & Henrick, 2007).

**Figure 1.**
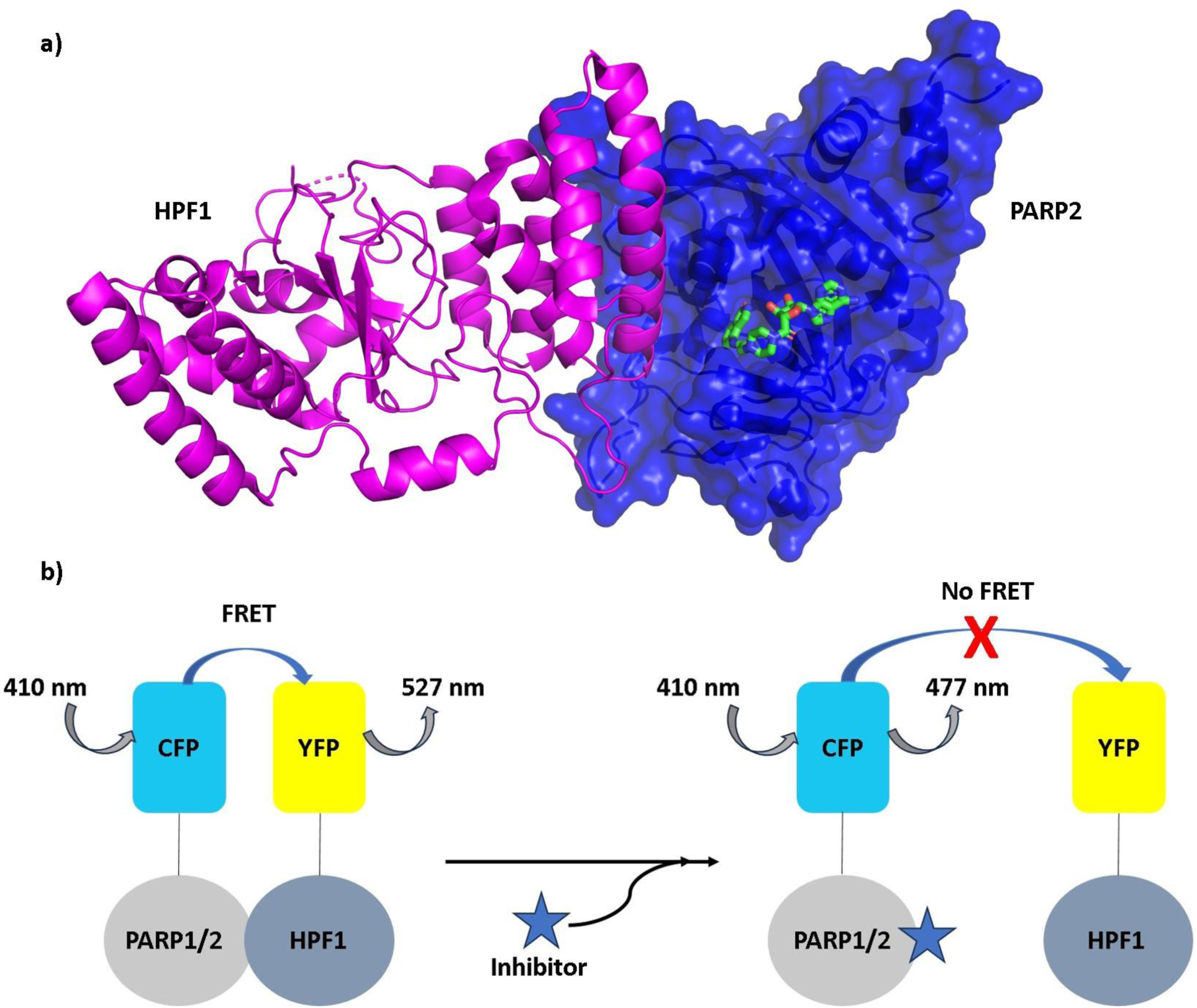
HPF1-PARP interaction. **a**) HPF1 (magenta) and PARP2 (blue) interaction interface. NAD^+^ analog EB-47 (green) is bound to the PARP2 catalytic site (PDB id: 6TX3) (Suskiewicz *et al*, 2020), **b**) HPF1-PARP FRET pair interaction (left panel) and inhibition of HPF1-PARP interaction by inhibitor resulting in loss of FRET (right panel).

In order to generate a signal that can be used in high-throughput mode to discover protein-protein interaction inhibitors as potential starting points for drug development, we decided to label the proteins with fluorophores to develop a FRET assay. We expressed fusion proteins where mCerulean (CFP) is attached to the N-terminus of the PARP catalytic domain and mCitrine (YFP) is attached to the N-terminus of HPF1 as then the tags would not interfere with the complex formation based on the structure. When there is an interaction between labeled HPF1 and PARP1/2 then YFP and CFP come in close proximity and upon excitation of CFP there is an energy transfer from CFP to YFP resulting in FRET signal (**Fig. 1b**). Protein-protein interaction inhibitor would result in a loss of the FRET signal. We optimized the assay for screening of inhibitors of PARP2-HPF1 interaction and showed that the homogeneous assay is robust and can be run on a 384-well microplates in high-throughput screening campaigns. During the assay validation, we discovered two compounds Dimethylacrylshikonin and Alkannin as inhibitors of the protein-protein interaction.

## Results

### Assay setup and FRET measurement

To measure the interaction between PARP1/2 and HPF1 and to develop a high throughput method for inhibitor screening we used a FRET-based approach. We cloned constructs of PARP1 and PARP2 catalytic domains fused with CFP at the N-terminus and HPF1 fused with YFP at the N-terminus. Purified CFP-PARP1/2 and YFP-HPF1 were mixed, and FRET was measured by excitation of CFP at 410 nm wavelength and measuring YFP fluorescence intensity (527 nm) and CFP fluorescence intensity (477 nm). When CFP and YFP interact, the CFP emission at 477 nm decreases while YFP emission at 527 nm increases. The ratio of emission intensities of 527 nm and 477 nm, rFRET, was used throughout the work as a measure of binding interaction when comparing it with controls. As the yields for produced CFP-PARP2 were higher than those for CFP-PARP1 (25 mg/l and 10 mg/l, respectively), we decided to use CFP-PARP2 for optimization and for screening. For the interaction between HPF1 and PARP2, buffer conditions i.e., pH, buffering reagent, buffer and salt concentration, additives were optimized to get a robust and reproducible signal (**Fig. 2**).

**Figure 2.**
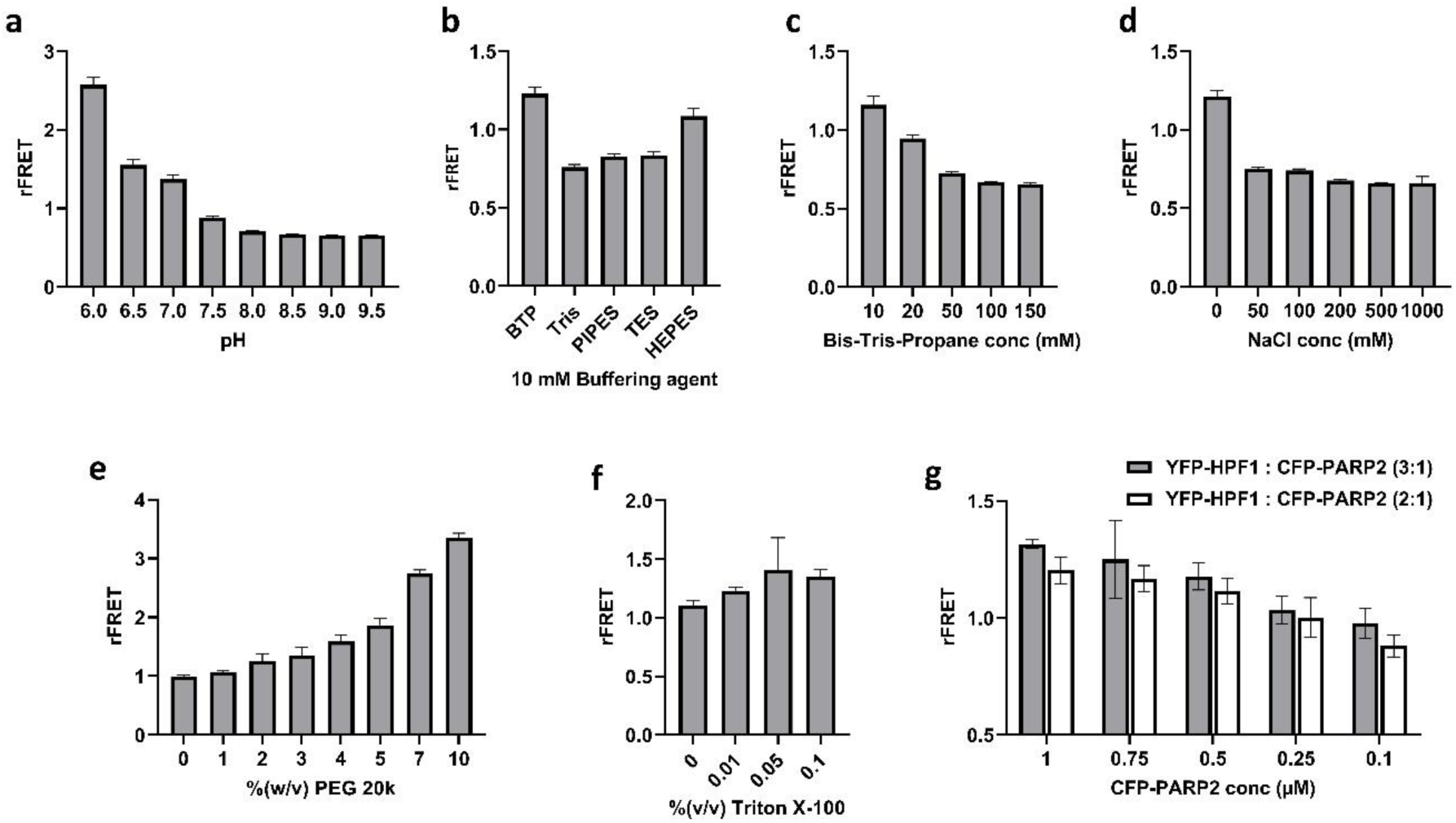
Buffer and protein concentration optimization. **a**) Buffer pH optimization in 10 mM Bis-Tris-Propane. **b**) Buffering reagent optimization with 10 mM concentration at pH 7.0. **c)** Bis-Tris-Propane (BTP) concentration optimization at pH 7.0. **d)** NaCl concentration optimization in 10 mM BTP at pH 7.0. **e)** %(w/v) PEG 20k optimization in 10 mM BTP at pH 7.0. **f)** Detergent Triton X-100 %(v/v) optimization in 10 mM BTP at pH 7.0 In panels a-f 5 µM YFP-HPF and 1 µM CFP-PARP2 were used in reactions. **g)** YFP-HPF1 and CFP-PARP2 protein concentration optimization in FRET buffer (10 mM BTP, 0.01% Triton X-100, 3% PEG 20k, pH 7.0) where x-axis is representing CFP-PARP2 concentrations (µM) and YFP-HPF1 was mixed in 2:1 (white bars) and 3:1 (grey bars) ratio to CFP-PARP2 concentration. All the buffer and protein concentration optimization were performed in 20 µl volume in 384 well plates and the data shown are mean with standard deviations of 4 replicates.

First the effect of pH on rFRET pair (5 µM YFP-HPF1 and 1 µM CFP-PARP2) was tested using 10 mM Bis-Tris-Propane (BTP), which has pH ranges from 6.0 to 9.5 (**Fig. 2a**). At lower pH at 6.0, YFP-HPF1 (5 µM) and CFP-PARP2 (1 µM) FRET pairs showed maximum signal but it was evident that at pH at 6.0 and 6.5 proteins were aggregating in the well and this was interpreted as the source of the higher rFRET signal. Increasing pH resulted in diminishing signal, while at neutral pH 7.0 the rFRET signal was present and there were no signs of protein aggregation in the well. Therefore, we decided to further optimize the buffer conditions at pH 7.0, which is also close to physiological pH and therefore the ionization state of the potential inhibitors will be reasonable. After optimizing pH, 10 mM buffering reagents at pH 7.0 were tested for the FRET pair and since the highest signal was observed with BTP we decided to continue with it (**Fig. 2b**). Increasing concentrations of the buffering agent, however, decreased the signal (**Fig. 2c**). This could be an indication of sensitivity of the system to increasing salt concentrations and indeed, when we tested NaCl concentrations from no salt to 1 M we observed that even low salt concentration of 50 mM drastically reduced the rFRET signal (**Fig. 2d**). We therefore decided to use no salt in the buffer and tested the effect of Polyethylene Glycol 20,000 (PEG 20k) as a crowding agent. PEG has shown to enhance the rFRET signal and help in blocking adhesion the biomolecules to plastic walls (Liu *et al*, 2013). Increasing PEG concentrations resulted in an increase in the rFRET signal also in this case (**Fig. 2e**). At higher PEG concentration rFRET signal increased very rapidly to unusually high rFRET values based on our previous experience (Sowa *et al*, 2020, 2021) and we interpreted this to be due to unspecific interactions. We decided therefore to add only a small amount of 3% (w/v) PEG 20k to the FRET buffer. Triton X-100 was found to slightly increase the signal and as it could help in the assay setup including the meniscus formation in the experimental plate, we also added 0.01% (w/v) to the final FRET buffer (**Fig. 2f**). Based on the optimization, 10 mM BTP, 0.01% tritonX-100, 0.5 mM TCEP, 3% PEG 20k, pH 7.0, was selected as an optimal FRET buffer. We next wanted to test the effect of HPF1 PARP2 stoichiometry and whether we would be able to reduce the proteins concentration while maintaining a good enough signal for screening. Increase in the relative HPF1 concentration slightly increased the rFRET signal systematically as more complex would be present in the mixture (**Fig. 2g**). The difference was however small and to save protein and to keep the sensitivity of the assay reasonable we made a compromise and decided to use 400 nM CFP-PARP2 and 800 nM YFP-HPF1 for further studies.

### Signal and protein stability

In the initial part of optimization procedure, we studied the stability of the rFRET signal over time (**Fig. 3a**). We observed that the rFRET signal was increasing with time indicating non-steady state conditions that could be a result of instability of the proteins. This increase in signal over time would be a problem also in the context of a larger screening campaign as the dispensing and readouts need to be precisely timed. As the preparation of PARP1 and PARP2 without the autoinhibitory helical domain has been challenging we suspected that the protein constructs would not be stable in the used low salt buffer and measured the thermal stability using differential scanning fluorimetry (nanoDSF). It was evident that PARP2 started unfolding and did not show any clear transition in the FRET buffer at very low temperatures (Tm 25.6 **±** 0.15°C) (**Fig. 3b)**.

**Figure 3.**
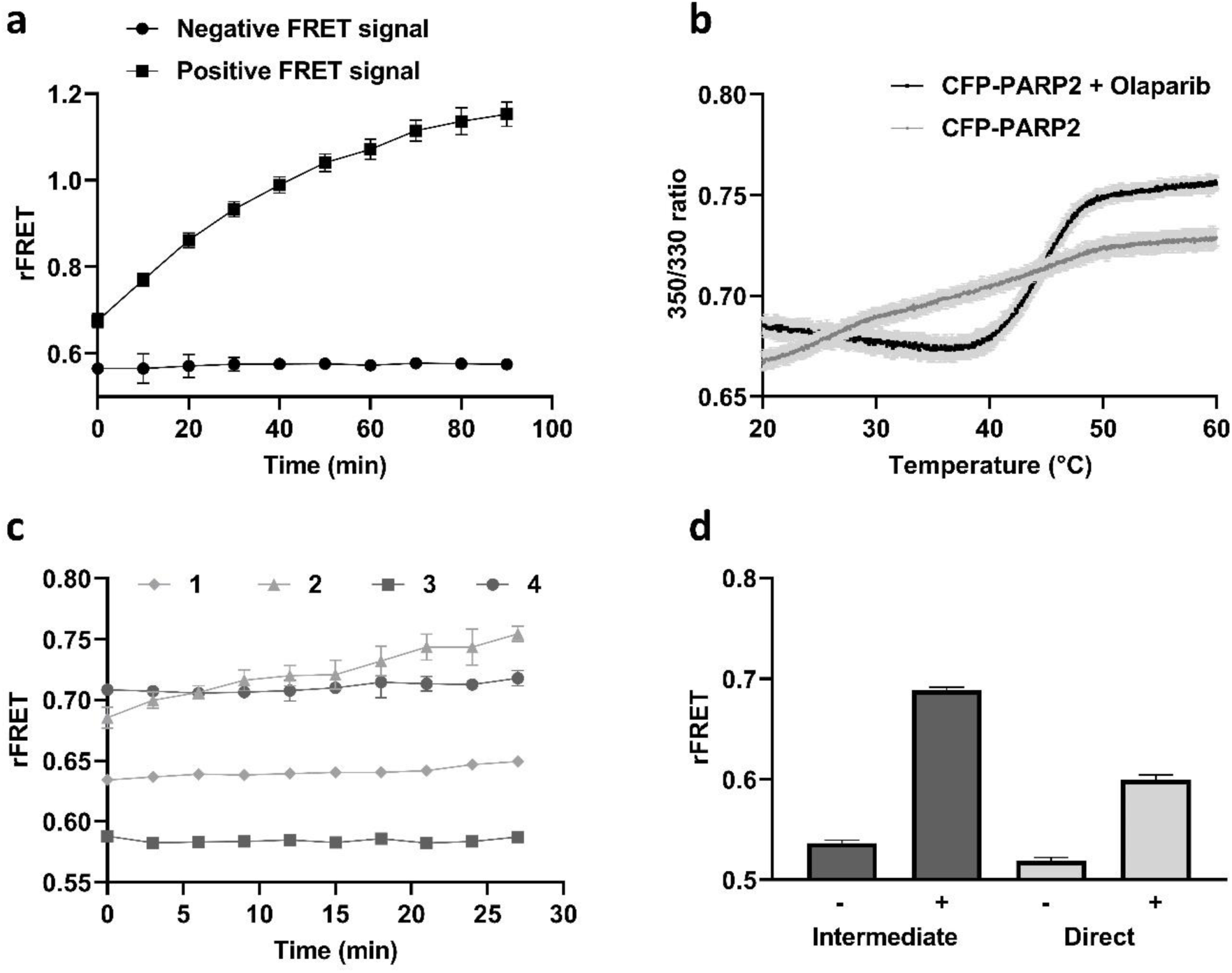
rFRET signal for CFP-PARP2 and YFP-HPF1. **a**) Time dependent monitoring of rFRET signal, where FRET pair concentration is 400 nM CFP-PARP2 and 800 nM YFP-HPF1. **b**) PARP2 catalytic domain (1 mg/ml) thermal stability without (grey) and with (black) Olaparib (100 µM). **c**) rFRET signal (400 nM CFP-PARP2 and 800 nM YFP-HPF1) monitoring with time in absence and presence of Olaparib [1 – negative rFRET signal without Olaparib, 2 – positive rFRET signal without Olaparib, 3 – negative rFRET pair signal with Olaparib, 4 – positive rFRET pair signal with Olaparib]; **d**) Effect of protein mixing on rFRET signal, [Dark grey bars: rFRET signal with intermediate dilution of protein (25 µM CFP-PARP2 and 50 µM YFP-HPF1) for no FRET (-) and FRET (+) reaction; Light grey bars: rFRET signal with mixing of protein (400 nM CFP-PARP2 and 800 nM YFP-HPF1) for no FRET (-) and FRET (+) reaction in the FRET buffer]. All the reactions were performed in 20 µl volume in 384 wells plate and the data shown are mean with standard deviations of 4 replicates.

The increasing rFRET signal and nanoDSF data, collectively suggested that the protein was not stable at room temperature and was aggregating with time potentially bringing the fluorophores to proximity. Buffer additives such as glycine, guanidium hydrochloride, polyoxyethylene lauryl ether (Brij-35) were tested to stabilize the CFP-PARP2 protein, but additives did not help in stabilizing the protein in FRET buffer (data not shown) (Bye *et al*, 2014; Schwarz *et al*, 2007). To stabilize the PARP2 fragment, 5-fold excess of PARP catalytic inhibitor Olaparib was added in the FRET buffer. Addition of Olaparib resulted in a sharp unfolding transition for PARP2 with higher Tm value (44.6 ± 0.07 °C) by making the protein stable at room temperature (**Fig. 3b**). For the fusion protein 2 transitions were expected as it contains 2 independent domains with different Tms. The thermal melting was run up to 60°C and Tm for CFP is 80°C, so a single transition was observed for PARP2 catalytic domain only. In the presence of Olaparib, CFP-PARP2 and YFP-HPF1 interaction was stable in the FRET assay and simultaneously confirmed that Olaparib was not interfering in the PARP2-HPF1 interaction and could be used in the interaction assay (**Fig. 3c**).

In the presence of Olaparib, 400 nM CFP-PARP2 and 800 nM YFP-HPF1 were mixed resulting in a stable signal within an experiment, but the rFRET values were not reproducible in different experiments. This could indicate variation in complex formation between experiments depending on the order of addition and salt concentration that comes from the independently purified proteins. After trial and error, FRET pair proteins were mixed first in higher concentration (25 µM CFP-PARP2, 50 µM YFP-HPF1) in the buffer containing excess Olaparib (100 µM) in a so called intermediate mixture. Final FRET mixture with 400 nM CFP-PARP2 and 800 nM YFP-HPF1 was then prepared by diluting the intermediate mixture in FRET buffer. Mixing of the proteins in high concentration allows formation of complex in saturating amounts before the dilution to the lower assay conditions. This procedure allowed us to get a stable, reproducible, and higher rFRET signal in comparison to a direct mixing to the final FRET conditions (**Fig. 3d**).

### Signal Verification

In the optimization phase we used rFRET as a direct readout for the interaction and secondary information like protein aggregation to judge whether the signal resulted from the interaction between the proteins. To confirm this, we next tested unlabeled HPF1 to compete with the FRET pair and expected that the competition would result in the loss of rFRET signal. Increasing concentration of unlabeled HPF1 in the reaction indeed decreased the rFRET signal in a concentration dependent manner, but it should be noted here that there is salt added also to the mixture coming from the unlabeled HPF1 protein stock (**Fig. 4a**).To rule out the effect of added salt, we repeated the experiment in the absence of HPF1 but using equivalent salt concentrations which resulted in higher IC_50_ (pIC_50_: 4.5 **±** 0.007, IC_50_: 29.07 µM) as compared to the competition by HPF1 (pIC_50_: 5.83 **±** 0.03, IC_50_: 1.46 µM). The 1.3-log unit difference in the apparent IC_50_ values clearly indicates that unlabeled HPF1 competes with the FRET pair (**Fig. 4a**). To further strengthen the analysis and to confirm the competition we also exchanged the HPF1 buffer. The desalted unlabeled HPF1 also showed concentration dependent lowering (pIC: 4.62, IC_50_: 24.0 µM) of the rFRET signal (**Fig. 4b**). We however observed aggregation of HPF1 in no salt condition at high concentrations and therefore the active unlabeled HPF1 protein concentration in the reaction was likely lower than the calculated total concentration while the competition was still evident. These results confirmed that unlabeled HPF1 (without salt) competed with the CFP-PARP2 and YFP-HPF1 FRET pair and validated the interaction between HPF1 and PARP2 in the FRET reaction.

**Figure 4.**
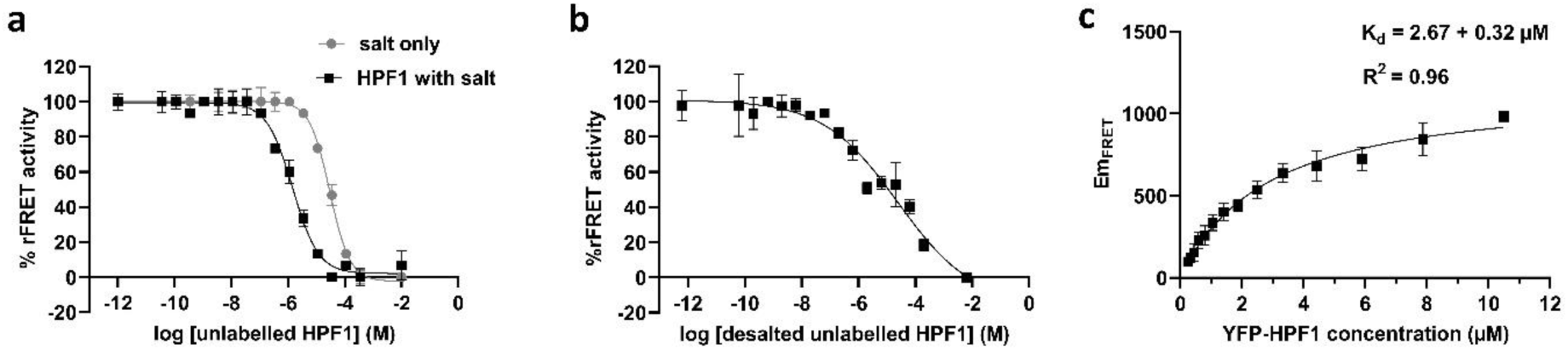
CFP-PARP2 and YFP-HPF1 FRET pair signal competition with unlabeled HPF1 and determination of dissociation constant. **a**) FRET pair competition with unlabeled HPF1 containing salt (black), equivalent salt only (grey). **b**) FRET pair competition with desalted unlabeled HPF1. **c**) FRET-based determination (Kd) for CFP-PARP2 with YFP-HPF1 interaction. The FRET fluorescence emissions (Em_FRET_) were determined as described by Song et al.(Song *et al*, 2012). All the reactions were performed in 20 µl volume in 384 well plates and the data shown are mean with standard deviations of 4 replicates.

### Dissociation constant for PARP2-HPF1 interaction

The binding affinity of PARP2-HPF1 was measured using FRET according to the method described by Song et. al **(Song *et al*, 2012)**. An increasing concentration of YFP-HPF1 (250 nM to 10.5 µM) was mixed with a constant concentration of CFP-PARP2 (400 nM) to saturate the CFP with YFP signal. Here we did not consider effect of salt as it is apparently affecting in low protein concentrations and at concentration >>Kd the complex is formed and can be even crystallized. Based on the raw fluorescence data, the FRET emission (Em_FRET_) was calculated, and the saturation curve showed Kd of 2.67 **±** 0.32 µM. This is in line with the values and estimates from the literature 3 µM for full length wild type PARP2 **(Gaullier *et al*, 2020)** and indicates that in the preparation of the FRET mixture the complex is formed in the intermediate mixing (25 µM CFP-PARP2, 50 µM YFP-HPF1) and this is then partially dissociated when diluted to the final FRET buffer (400 nM CFP-PARP2 and 800 nM YFP-HPF1).

### Signal validation for screening

Once the CFP-PARP2 and YFP-HPF1 interaction was verified, the rFRET signal performance was validated by repeating the experiments on different days using different plates. The experiments were performed in 10 µl and 20 µl reaction volumes in 384-well plates, which contained 176 replicates of FRET pair (positive control) and 176 replicates of FRET pair with 1 M NaCl (negative control) (**Fig. 5**). The Zʹ-factors for 10 µl and 20 µl FRET reaction volumes were determined in a range of 0.85 to 0.92 and 0.75 to 0.88, respectively (**Table S1 and S2**). The Z’ factor value above 0.5 indicates the separation between negative and positive control is large, and assay with Z’ value above 0.5 is considered as excellent **(Zhang *et al*, 1999)**. The statistical analysis suggests that even in a low assay volume of only 10 µl the quality of the signal was consistent, and the assay can be used for the screening of inhibitors.

**Figure 5.**
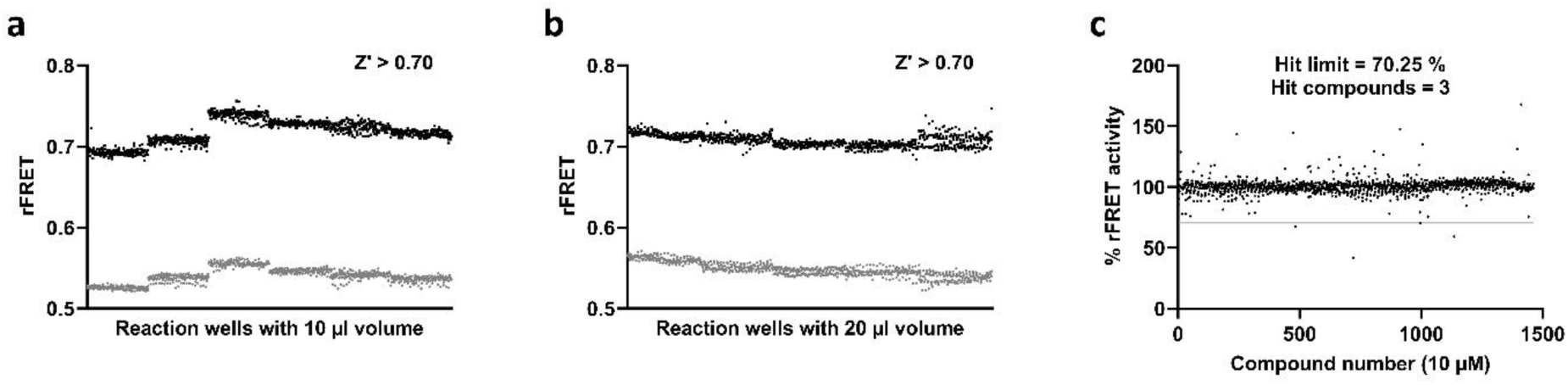
Signal validation and validatory screening in 384-well plates. **a**) Signal validation in 10 µl reaction volume. **b**) Signal validation in 20 µl reaction volume. **c**) Validatory screening using Targetmol compound library with 10 µM compound in 10 µl final reaction volume.

### Validatory screening

To validate the assay for inhibitor screening, Targetmol inhibitor compound library (1832 compounds), which contains biologically active compounds, was screened (**Fig. 5c**). For the screening, 10 nl compounds (10 µM concentration in reaction) and 10 µl FRET pair protein mixture (400 nM CFP-PARP2 and 400 nM YFP-HPF1) was dispensed and measured. In presence of inhibitors, an additional excitation wavelength 430 nm was also used for the FRET measurement to rule out spurious effects due to compound fluorescence. This is based on the idea that for true protein-protein interaction inhibitors the inhibition at different excitation wavelengths is expected to be similar. On the other hand, artifacts due to small molecules fluorescence are expected to have different effects at different excitation wavelengths since they usually have sharper emission and excitation spectra. To analyze inhibition data, multiple filters were applied to rule out non-specific FRET signal coming from intrinsic fluorescence of individual compounds. Raw fluorescence signal with more than 20% fluorescence value (>1.2* fluorescence at 527 nm; <1.2*fluorescence at 477 nm) from the positive control FRET pair fluorescence signal, were considered as outlier and were not included for further analysis (**Table S3**). The % rFRET activity was calculated for each compound based on the controls. The hit limit for library compounds was set to 70.25% activity (5* standard deviations from mean) (**Table S3**). In the screening, a total of 3 compounds Dimethylacrylshikonin (CAS no. 24502-79-2), Alkannin (CAS no. 517-88-4), Crystal violet (CAS no. 548-62-9) were identified as hits for PARP2-HPF1 interaction.

### Hit validation

To validate the hit compounds from CFP-PARP2 and YFP-HPF1 FRET pair screening, the binding of the compounds (Dimethylacrylshikonin, Alkannin, Crystal violet) with PARP2 and HPF1 were measured using nanoDSF (**Fig. S2**). When compound binds to the protein it results in the increase in the melting temperature of the protein. The Tm values for PARP2 and HPF1 in absence of hit compound were 44.2 ± 0.10 °C and 46.8 ± 0.02 °C, respectively. In the presence of hit compounds Dimethylacrylshikonin, Alkannin, and Crystal violet, the Tm value of PARP2 were measured as 46.0 ± 0.4 °C, 46.0 ± 0.2 °C, and 40.0 ± 0.5 °C, respectively (**Fig. S2 a-c**). The increase in Tm in presence of Dimethylacrylshikonin and Alkannin, suggested their binding to PARP2 catalytic domain, but Crystal violet destabilizes the protein. The compounds did not cause a thermal shift for HPF1.

PARP2 catalytic domain has been crystallized with HPF1 and the catalytic domain of PARP1 and PARP2 shares a similar tertiary structures (Suskiewicz *et al*, 2020). PARP1 is also interacting with HPF1 approximately with the same interface as PARP2 and forming a joint active site with HPF1 (**Fig. S1**). To study also the interaction between PARP1 catalytic domain and HPF1, CFP-PARP1 was mixed with YFP-HPF1 in 1:2 concentration ratio in optimized buffer for PARP2-HPF1 FRET pair (**Fig. 6a**). The FRET was measured using the same parameters as used for CFP-PARP2 and YFP-HPF1 FRET pair and a stable FRET signal was observed (**Fig.6b**). This allowed us to measure potencies of compounds for both, PARP1-HPF1 and PARP2-HPF1, interactions.

**Figure 6.**
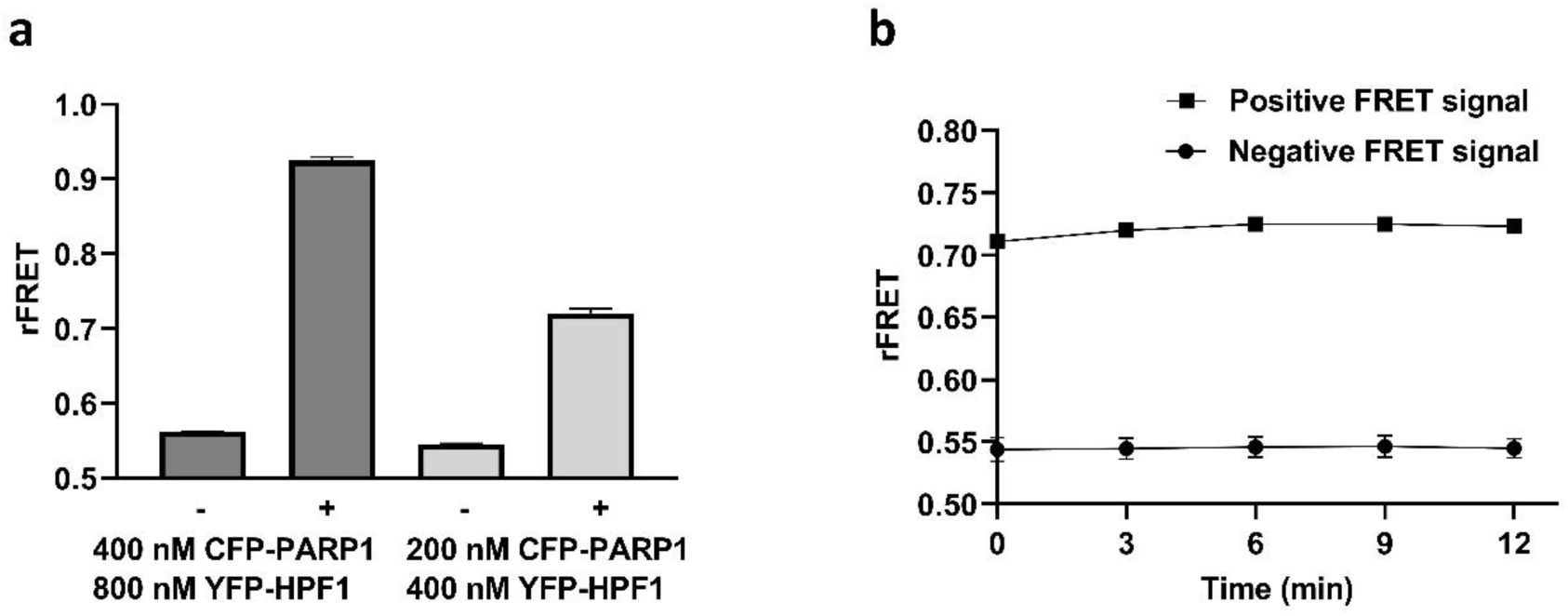
rFRET signal for **CFP-**PARP1 and YFP-HPF1 FRET pair. **a**) rFRET signal from control (-) and FRET pair (+) at different protein concentrations of CFP**-**PARP1 and YFP-HPF1. **b**) Time dependent monitoring of rFRET signal, with 200 nM CFP-PARP1 and 400 nM YFP-HPF1 protein concentration. All the reactions were performed in 10 µl volume in 384 well plates and the data shown is mean ± standard deviation with 4 replicates.

To confirm the inhibition and to identify the potency of Dimethylacrylshikonin and Alkannin against PARP1/2 and HPF1 interaction, their IC_50_ values were measured. The IC_50_ values for Dimethylacrylshikonin, Alkannin against PARP1-HPF1 complex were 29.30 µM (pIC_50_: 4.53 ± 0.40) and 16.23 µM (pIC_50_: 4.80 ± 0.20), respectively (**Fig. 7a, b**) while IC_50_ values against PARP2-HPF1 complex were 15.73 µM (pIC_50_: 4.80 ± 0.17) and 21.20 µM (pIC_50_: 4.67 ± 0.06), respectively (**Fig. 7c, d)**. Based on the binding of compounds to PARP catalytic domain and IC_50_ measurements, Dimethylacrylshikonin and Alkannin were identified as first described inhibitors of the PARP1/2-HPF1 interaction.

**Figure 7.**
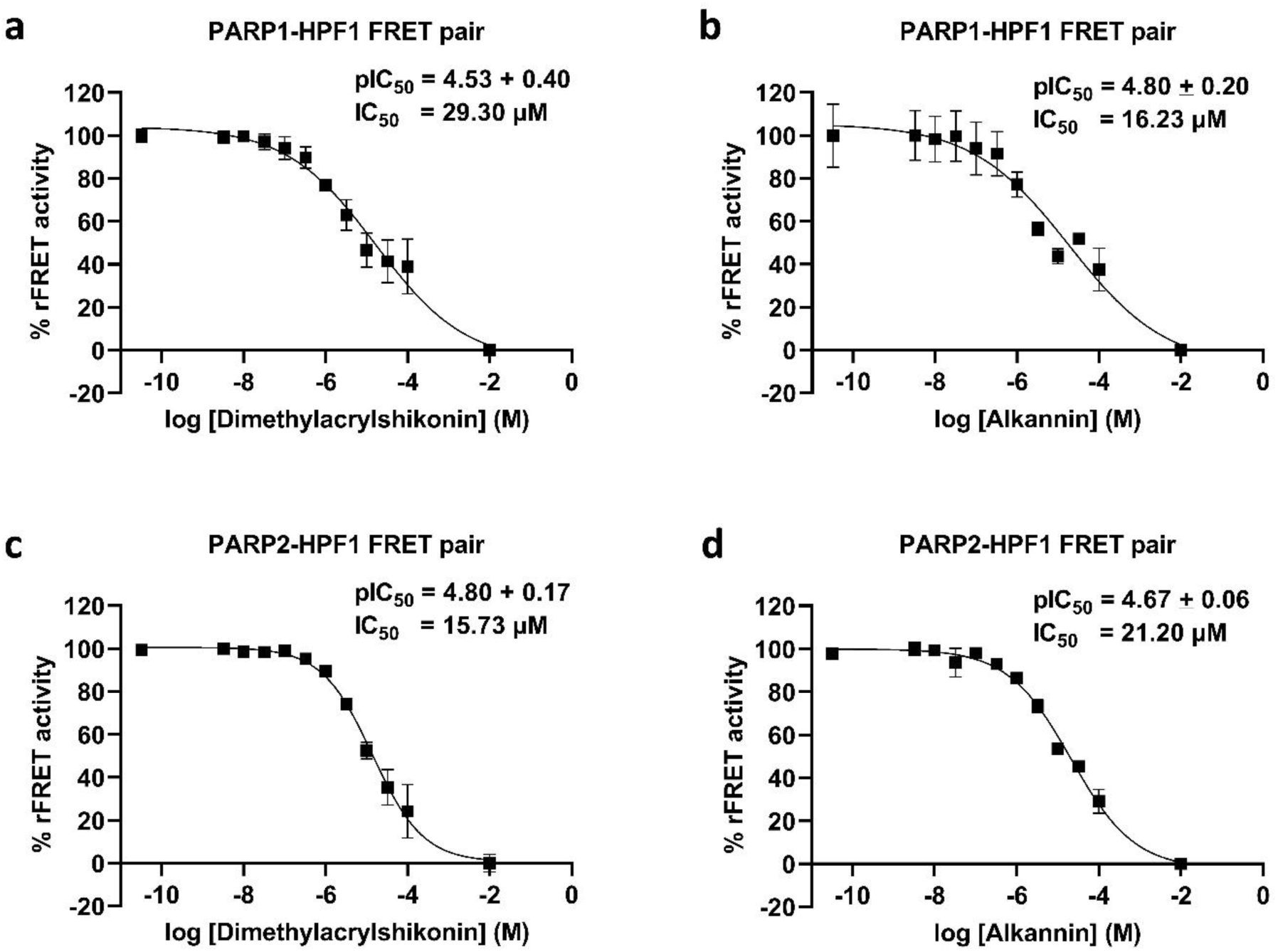
Example IC_50_ curves for Inhibition of PARP-HPF1 interaction by increasing concentration of inhibitor compounds. PARP1-HPF1 interaction inhibition by **a)** Dimethylacrylshikonin **b)** Alkannin. PARP2 -HPF1 inhibition by **c)** Dimethylacrylshikonin **d)** Alkannin.

## Discussion

Current PARP inhibitors target the catalytic site and compete with NAD^+^. The discovery that the ADP-ribosylation is shifted to serines in the context of DNA damage repair by HPF1 makes the interference an alternative and unique way to inhibit PARP activity (Bonfiglio *et al*, 2017; Suskiewicz *et al*, 2020; Sun *et al*, 2021). Protein–protein interactions are known to be challenging to target and so far there are no inhibitors reported for the PARP-HPF1 interaction. In this study, we have developed a FRET-based high-throughput screening assay that allows direct monitoring of HPF1-PARP interaction. The assay can be used for screening of small inhibitor molecules against PARP-HPF1 interaction. The proteins are expressed as fusion proteins with fluorescent proteins and when PARP interacts with HPF1 the FRET signal can be monitored using a common plate reader fluorimeter. We validated that the signal results from the specific interaction by competing it out with unlabeled HPF1. The signal and interaction in nanomolar concentrations is sensitive to salt making it convenient to use just high NaCl concentration for the control in lack of known inhibitors. Extensive efforts in optimization resulted in an assay which showed consistently a stable and robust signal (Z’> 0.7) and can be run with automation in low 10 µl volumes in 384-well plates. The assay requires >100 nM protein concentrations, but, as specialized reagents are not needed and the recombinant fusion proteins are produced in *E. coli*, the assay is cost effective and can be applied to large screening campaigns.

Recently, there was a study to find inhibitors against HPF1-PARP1 complex, where the interference of compounds between HPF1 and PARP1 was indirectly monitored by measuring the retention time of PARP1 on DNA (Kellett *et al*, 2023). This fluorescence polarization assay also identified catalytic PARP inhibitors and therefore the mechanism of the identified compounds may not be evident. In contrast, in the FRET-assay we describe here we measure directly the interaction between HPF1 and PARP and the hits, if not interfering with the signal or destabilizing proteins like crystal violet, are expected to act as protein-protein interaction inhibitors. The assay works even in the presence of a catalytic clinically used inhibitor and therefore selects the inhibitors mechanistically and avoids compounds only binding to the catalytic site unless the hit compounds extend from the catalytic site to the HPF1 binding interface. This feature extends the utility of the assay to test the mechanism of inhibitors identified by other means (Choo *et al*, 2021; Kellett *et al*, 2023).

During the validatory screening we discovered two hit compounds Dimethylacrylshikonin and Alkannin. These compounds stabilized PARP2 indicating direct binding and had µM potencies for both PARP1-HPF1 and PARP2-HPF1 interaction. The screening method will hopefully facilitate the discovery of potent inhibitors against HPF1-PARP1/2 complexes and help to develop effective therapeutic molecules for cancer with a unique mechanism.

## Materials and Methods

### Cloning

Inserts of PARP1/2 catalytic domain and full length HPF1 were prepared using gene specific primer in PCR. To make expression construct, PARP1 catalytic domain [T661-F676+ GSGSGSGG+(D788-W1014)] (Dawicki-McKenna *et al*, 2015) or PARP2 catalytic domain (without regulatory helical domain) were cloned into pNIC-CFP plasmid (<u>addgene # 173083)</u> and HPF1 was cloned into pNIC-YFP plasmid (addgene # 173080) using SLIC cloning method (Jeong *et al*, 2012). The pNIC-CFP or pNIC-YFP plasmids were linearized in PCR using site specific primers. 100 ng linearized plasmid was mixed in 1:3 molar ratio with gene PCR products and incubated with T4 DNA polymerase for 2-3 minutes at room temperature. The mixture was transformed into *E. coli* strain NEB5α cells and colonies were grown at 37 ^°^C overnight on LB agar media plates containing, Kanamycin as antibiotic, 10 mM Benzamide, and 5% sucrose for SacB-based negative selection (Hynes *et al*, 1989; Reyrat *et al*, 1998). Plasmid of CFP tagged PARP1 contains “His6 tag-CFP-TEV protease site-catalytic domain [T661-F676+GSGSGSGG+ (D788-W1014)]”, PARP2 construct contains “His6 tag-CFP-TEV protease site-catalytic domain (without regulatory helical domain)” and YFP tagged HPF1 contains His6 tag-CFP-TEV protease site followed by full length HPF1.Protein expression. Plasmid containing CFP-PARP2 and YFP-HPF1 were transformed into *E. coli* BL21(DE3) and CFP-PARP1 was transformed to *E. coli* Rosetta2 (DE3) cells for the protein expression and incubated at 37^0^C overnight. Transformed colonies were further inoculated in 500 ml autoinduction media which contains trace elements and supplied with 0.8% (w/v) glycerol, 50 μg/ml kanamycin antibiotic. PARP inhibitors 10 mM Benzamide and 2 mM 3-Amino Benzamide were added to the culture medium of CFP-PARP2 and CFP-PARP1, respectively (Banasik *et al*, 1992). Cultures were incubated at 200 rpm at 37 °C until an OD600 of 1.2, and then incubated for 18 h at 16 °C. Next day, cells were harvested by centrifugation at 4,200×g for 45 min at 4 °C. Pellets were resuspended in lysis buffer (50 mM HEPES, 350 mM NaCl, 10 mM imidazole, 10% (w/v) glycerol, pH 7.5) and 100 µM pefablock [4-(2-Aminoethyl) benzenesulfonyl fluoride hydrochloride] protease inhibitor was added in it. Resuspended cells were flash frozen in liquid nitrogen and stored at -20 °C.

Protein purification. All the constructs (CFP-PARP1, CFP-PARP2, YFP-HPF1) were purified using two step purification i.e., immobilized metal affinity chromatography (IMAC) and size exclusion chromatography (SEC). Harvested cells were thawed, DNase (2 µg/ml) was added, and cells were lysed by sonication. Lysate was centrifuged at 39000xg RCF, for 1hr at 4°C, supernatant was filtered and loaded on lysis buffer equilibrated, Ni-NTA column. The column was washed with 3 column volumes wash buffer 1 (30 mM HEPES, 1 M NaCl, 10% glycerol, 0.5 TCEP, 10 mM imidazole, pH 7.5) and wash buffer 2 (30 mM HEPES, 500 mM NaCl, 10% glycerol, 0.5 mM TCEP, 50 mM imidazole, pH 7.5). Protein was eluted in elution buffer (30 mM HEPES [pH7.5], 500 mM NaCl, 10% glycerol, 0.5 mM TCEP, 250 mM Imidazole, pH 7.5). The eluted protein was loaded on pre-equilibrated (30 mM HEPES, 500 mM NaCl, 10% glycerol, 0.5 mM TCEP, pH 7.5) 16/600 superdex-75 size-exclusion chromatography column. Purified proteins were concentrated using 30 kDa membrane filter, aliquoted in small volumes, flash-frozen in liquid nitrogen and stored at -70 °C. The yield of proteins for YFP-HPF1, CFP-PARP2 and CFP-PARP1 were approximately 70 mg/liter, 25 mg/liter and 10 mg/liter, respectively. For YFP-HPF1 and CFP-PARP1/2 all steps of protein purification were performed at 4 °C in a cold room.

### FRET measurement

FRET signal measurements were performed using Tecan Spark multimode plate reader. Measurement was done using 384-well plate (proxiplate-384 F plus from Perkin Elmer). CFP was excited using monochromator at 410 nm (20 nm bandwidth) excitation wavelength, emission was recorded at emission wavelengths of 477 (10 nm bandwidth) and 527 nm (10 nm bandwidth). Other parameters such as number of flashes 50, Z position 20500, signal integration time 40 µs, settle time 10 ms, manual gain 90 were optimized and used for the FRET measurement. To calculate the ratiometric FRET (rFRET) value, fluorescence intensity at 527 nm was divided by fluorescence intensity at 477 nm. All the rFRET measurements were done in optimized buffer 10 mM Bis–Tris-Propane, 0.5 mM TCEP, 3% (w/v) PEG 20k, and 0.01%(v/v) TritonX-100, pH 7.5, unless otherwise stated in negative control samples, 1 M NaCl was added.

### Protein Stability

Protein stability was tested using nanoDSF (differential scanning fluorimetry), where protein unfolding is monitored by tryptophan fluorescence. Different protein stabilizing regents, additives and PARP inhibitors were tested with CFP-PARP2 and YFP-HPF1 to see the effect on their melting temperatures (Tm values) and stability of CFP-PARP2 protein at room temperature.

### Competition assay

FRET Signal verification. Competition assay was performed with an increasing concentration of unlabeled HPF1 and NaCl. In this experiment, the increasing log concentration of unlabeled HPF1 or NaCl was mixed with CFP-PARP2 (400 nM) and YFP-HPF1 (800 nM) FRET pair. FRET pair with 1 M NaCl and without unlabeled HPF1 were used as lowest and highest FRET signal controls, respectively. Data were measured in 4 replicates with 20 µl reaction volume in 384-well microplate. Statistics of the data and graph were analyzed using GraphPad Prism 9.0 using nonlinear regression analysis and data was plotted as inhibitor concentration v/s rFRET signal.

### Determination of dissociation constant

To determine the dissociation constants of the FRET pair CFP-PARP2 and YFP-HPF1, constant CFP-PARP2 (400 nM) concentrations were mixed with increasing concentration of YFP-HPF1. First CFP-PARP2 and HPF1 were mixed in high concentration with increasing concentration of YFP-HPF1 in presence of Olaparib. The high concentration of YFP-HPF1 adds salt to the reaction from stock. This additional salt was counter balanced in all the reactions by backfilling of salt in buffer. The final reaction concentrations were achieved by mixing the dilutions to the buffer. The emission fluorescence was measured at 477 nm (10 nm bandwidth), 527 nm (10 nm bandwidth) by excitation at 430 nm (20 nm bandwidth), and emission at 527 nm (10 nm bandwidth) by excitation at 477 nm (20 nm bandwidth) wavelength. All other parameters for fluorescence measurement were kept the same as described in FRET measurements. Reaction was monitored in 10 µl volume with 4 replicates in 384-well plates. For the confidence in the Kd value, the experiments were performed 3 times, independently. Based on the raw fluorescence data, the FRET emission (Em_FRET_) was calculated using a formula “Em_FRET_ = F_527 at 430 excitation_ – α (F_477 at 430 excitation_) – β (F_527 at 477 excitation_)”. The fluorescence correction factor for CFP (α) and for YFP (β) were calculated as 0.477 and 0.058, respectively, using pure CFP and YFP constructs. The calculated Em_FRET_ intensities were fitted in GraphPad Prism 9.0 and Kd value was calculated using nonlinear regression method with a the equation “Y=EmFRETmax- ((2*EmFRETmax*Kd)/(X-A+Kd+sqrt(sqr(X-A-Kd)+(4*Kd*X))))” as described by Song et al **(Song *et al*, 2012)**.

### Assay signal validation

The assay signal verification was done with 10 µl and 20 µl reaction volume. To verify the signal, FRET pair protein mixture (400 nM CFP-PARP2, 800 nM YFP-HPF1) in optimized buffer (10 mM Bis–Tris-Propane, 0.5 mM TCEP, 3% (w/v) PEG 20k, 0.01%(v/v) TritonX-100, pH 7.5 and 1 µM Olaparib) with and without 1 M NaCl (192 replicates each condition) was dispensed to the 384-well plate using liquid dispenser robot (Formulatrix Mantis). This experiment was repeated for different days and with different plates on the same day. Data was analyzed in GraphPad Prism 9.0 using nonlinear regression curve fit and Z’-factor was calculated for the confidence in the assay **(Zhang *et al*, 1999)**.

### Validatory screening

For the validatory screening, 10nl of the compounds from Targetmol inhibitor compound library (10 mM stock) were transferred to the 384-well plates using small volume dispensing robot (Echo 650, Beckmann Coulter). 10 µl reaction volume of FRET pair protein mixture (400 nM CFP-PARP2, 800 nM YFP-HPF1) in optimized buffer (10 mM Bis–Tris-Propane, 0.5 mM TCEP, 3% (w/v) PEG 20k, 0.01%(v/v) TritonX-100, pH 7.0 and 1 µM olaparib) was dispensed in wells containing compounds using liquid dispenser robot Formulatrix Mantis. Fluorescence intensities at 477 nm and 527 nm were measured by exciting the reaction sequentially at 410 nm and 430 nm wavelengths. To analyze inhibition data from libraries, filters were applied to the remove non-specific FRET signal coming from intrinsic fluorescence of individual compounds. The %rFRET activity was calculated from rFRET signal (Emission_527_ / Emission_477_) using formula “%rFRET = [{(rFRET_reaction_ – rFRET _negative control_) / (rFRET_positive control_ – rFRET_negative control_)}*100]”. The statistical analysis of inhibition was performed in GraphPad Prism 9.0 using nonlinear regression curve fit.

### Hit Validation

To study the binding study of hit compounds from screening was done using nanoDSF. For the experiment, unlabelled full length HPF1 and PARP2 catalytic domain (without regulatory domain) with 1.0 mg/ml concentration were mixed in assay buffer (10 mM Bis–Tris-Propane, 0.5 mM TCEP, 3% (w/v) PEG 20k, and 0.01 %(v/v) Triton X-100, pH 7.5). Additionally, PARP2 buffer was supplied with excess of PARP catalytic inhibitor Olaparib (100µM PARP conc). Screened hit compounds (100 µM) from the FRET assay were added to the reaction mixture. The thermal melting scans were recorded in 3 replicates in three independent experiments from 20 °C to 60 °C with the temperature increase of 1 °C / min using Prometheus nanoDSF. The data was analyzed, and graphs were plotted using GraphPad Prism 9.0.

To validate the inhibition of FRET pair by hit compounds, the 50% inhibition concentration (IC_50_) values inhibitors against PARP1/2-HPF1 interaction, were measured using dilution series (100 µM to 3.25 nM) of compounds. Data was plotted against log inhibitor concentrations vs rFRET signal, analyzed in GraphPad Prism 9.0 using nonlinear regression analysis and IC_50_ values for compounds were calculated.

## Acknowledgements

The use of the facilities of the Biocenter Oulu Structural Biology core facility, member of Biocenter Finland, Instruct-ERIC Centre Finland and FINStruct, is gratefully acknowledged. The research was supported by the Research Council of Finland (Grant No. 347026) and by the Biocenter Oulu spearhead project funding.

## Author contributions

S.S.D. carried out most of the experiments and compiled the manuscript. A.G.-P. tested the initial assay setup. L.L. conceived the study and L.L. and A.G.-P. supervised the work. All authors contributed to the final version of the manuscript.

