## Supplementary figures and tables for "High-throughput screening assay for PARP-HPF1 interaction inhibitors to affect DNA damage repair"

### CONTENT

**Table S1 & S2.** Statistics of assay validation.

**Table S3.** Results of the compound screening.

**Figure S1.** Structural alignment of PARP1 catalytic domain with PARP2-HPF1 complex.

**Figure S2**. Thermal melting of PARP2 and HPF1 with inhibitors using nanoDSF.

**Table S1.** Statistics of assay validation at 10 µl volume.

| Day and plate number | Minimum Signal (control) | | | Maximum Signal (FRET pair) | | | S/N | Z' |
|---|---|---|---|---|---|---|---|---|
| | Average | SD | % CV | Average | SD | % CV | | |
| Day 1, plate 1 | 0.53 | 0.002 | 0.33 | 0.69 | 0.004 | 0.59 | 96.07 | 0.89 |
| Day 1, plate 2 | 0.54 | 0.004 | 0.71 | 0.71 | 0.004 | 0.59 | 44.51 | 0.86 |
| Day 2, plate 1 | 0.56 | 0.003 | 0.50 | 0.73 | 0.005 | 0.70 | 66.49 | 0.87 |
| Day 2, plate 2 | 0.55 | 0.002 | 0.43 | 0.73 | 0.003 | 0.37 | 77.26 | 0.92 |
| Day 3, plate 1 | 0.54 | 0.004 | 0.72 | 0.72 | 0.005 | 0.73 | 46.87 | 0.85 |
| Day 3, plate 2 | 0.54 | 0.003 | 0.64 | 0.71 | 0.003 | 0.46 | 52.78 | 0.89 |

**Table S2.** Statistics of assay validation at 20 µl volume.

| Day and plate number | Minimum Signal (control) | | | Maximum Signal (FRET pair) | | | S/N | Z' |
|---|---|---|---|---|---|---|---|---|
| | Average | SD | % CV | Average | SD | % CV | | |
| Day 1, plate 1 | 0.56 | 0.004 | 0.63 | 0.71 | 0.004 | 0.59 | 43.20 | 0.85 |
| Day 2, plate 1 | 0.55 | 0.004 | 0.69 | 0.71 | 0.005 | 0.70 | 42.12 | 0.84 |
| Day 2, plate 2 | 0.55 | 0.004 | 0.65 | 0.70 | 0.003 | 0.37 | 44.02 | 0.88 |
| Day 2, plate 3 | 0.54 | 0.004 | 0.78 | 0.70 | 0.004 | 0.53 | 36.61 | 0.85 |
| Day 3, plate 1 | 0.54 | 0.005 | 0.97 | 0.71 | 0.009 | 1.25 | 32.36 | 0.75 |

**Table S3.** Results of the compound screening.

| | |
|---|---|
| Number of compounds | 1832 |
| Filtered compounds (20% fluorescence filter) | 1462 |
| Mean of % activity | 100.2 |
| Std. Deviation | 6.08 |
| Hit limit (% activity) | 70.25 |
| Hit compounds | 3 |
| Selected after binding study (nano DSF) | 2 |

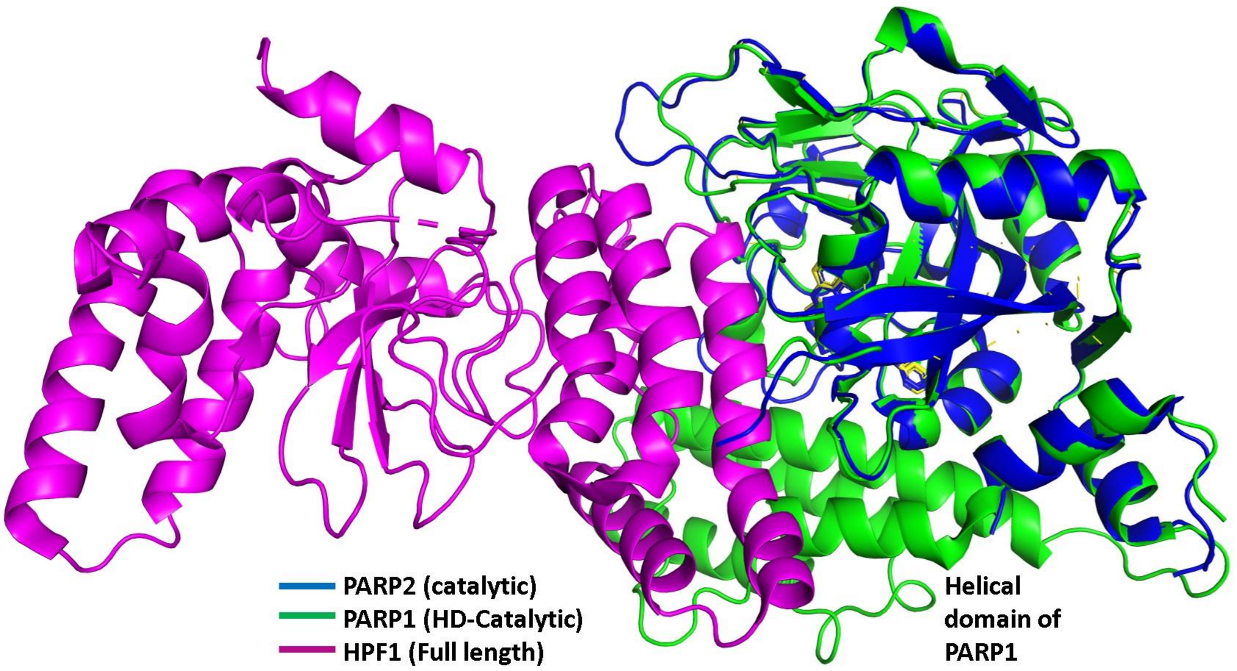


**Figure S1**. Structural alignment of PARP1 (HD-catalytic) (PDB id: 7AAB) with PARP2 (catalytic only)-HPF1 complex (PDB id: 6TX3). The catalytic domain of PARP1 (green) is superimposed with PARP2 catalytic domain interacting similarly with HPF1 as PARP2.

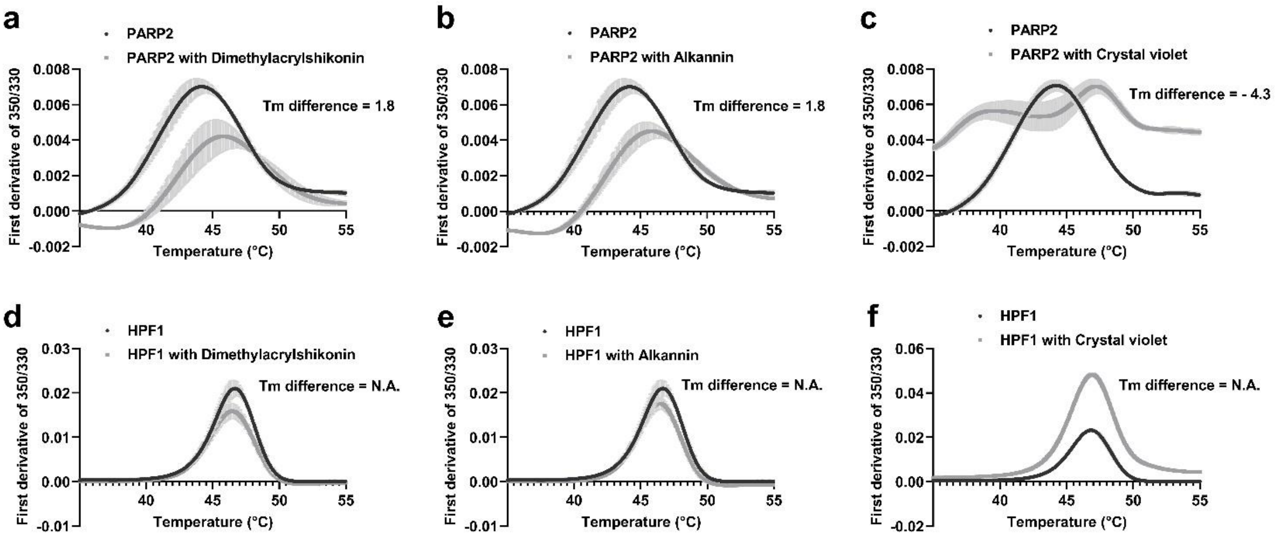


**Figure S2.** Thermal melting of protein (1 mg/ml) using nanoDSF showing the effect of hit compounds (100 μM) on thermal stability of (**a-c**) PARP2, (**d-f**) HPF1. The data shown are mean ± standard deviation from three independent measurements each with 3 internal replicates. [N.A.: not a significant shift in Tm]
